# The effects of luminal and trans-endothelial fluid flows on the extravasation and tissue invasion of tumor cells in a 3D *in vitro* microvascular platform

**DOI:** 10.1101/2020.09.23.309872

**Authors:** Cynthia Hajal, Lina Ibrahim, Jean Carlos Serrano, Giovanni S. Offeddu, Roger D. Kamm

**Author notes:** Correspondence to Roger D. Kamm, Department of Biological Engineering, Massachusetts Institute of Technology, NE47-318, Cambridge, MA, 02139, USA.

## Abstract

Throughout the process of metastatic dissemination, tumor cells are continuously subjected to mechanical forces resulting from complex fluid flows due to changes in pressures in their local microenvironments. While these forces have been associated with invasive phenotypes in 3D matrices, their role in key steps of the metastatic cascade, namely extravasation and subsequent interstitial migration, remains poorly understood. In this study, an *in vitro* model of the human microvasculature was employed to subject tumor cells to physiological luminal, trans-endothelial, and interstitial flows to evaluate their effects on those key steps of metastasis. Luminal flow promoted the extravasation potential of tumor cells, possibly as a result of their increased intravascular migration speed. Trans-endothelial flow increased the speed with which tumor cells transmigrated across the endothelium as well as their migration speed in the matrix following extravasation. In addition, tumor cells possessed a greater propensity to migrate in close proximity to the endothelium when subjected to physiological flows, which may promote the successful formation of metastatic foci. These results show important roles of fluid flow during extravasation and invasion, which could determine the local metastatic potential of tumor cells.

## INTRODUCTION

Metastatic seeding, which results in the dissemination of tumor cells (TCs) to secondary sites in the body, is the most common and lethal cancer complication^1^. To colonize distant organs, TCs must invade the surrounding tissue of the primary tumor, intravasate into the blood or lymph vasculature, travel to distant sites, extravasate from the circulation into the secondary organ, and survive, proliferate, and colonize the new microenvironment^2^. The latter steps involving the transmigration of TCs across the endothelium and their survival at the secondary site have been defined as the key rate-limiting steps of the metastatic cascade^3^, yet they remain poorly characterized due to experimental and imaging limitations^4,5^.

Cells and tissues constantly experience mechanical forces, such as those produced by fluid flow, particularly at the site of TC arrest in the metastatic organ. These fluid flows are vastly heterogeneous in the circulation and interstitium, due to varying hydrostatic and oncotic pressure gradients generated from differences in vessel geometry, endothelial permeability, and matrix fiber architecture^6–8^. In the circulation, TCs are subjected to fluid shear forces that affect their ability to adhere to the endothelial surface and alter their behavior through mechanically-activated signaling pathways. In addition to luminal flow, transmural or trans-endothelial (TE) flow occurs from the blood vessels into the surrounding matrix as interstitial flow, until its uptake by lymphatic vessels, which recirculates it to blood vessels^9,10^. These complex flow profiles in the circulation and surrounding tissues can result in significant biomechanical forces that have been shown to affect remodeling and angiogenesis^11^. More importantly, these varying biomechanical forces resulting from complex fluid flows may ultimately affect key steps of the metastatic cascade, playing a role in the ability of TCs to extravasate from the vasculature, migrate, and survive in the interstitium.

Previous studies into the role of hemodynamic and interstitial forces on TCs after they leave the primary site are sparse, although it is recognized that these forces play a role in determining migration characteristics^12–14^ and the metastatic potential of TCs^15–17^, ultimately affecting cancer patient outcomes^1^. Luminal flow and the shear stress it exerts, for example, can determine the arrest and initial endothelial adhesion of tumor cells^15,18^. However, *in vitro* studies investigating TC extravasation after such adhesion to the endothelium have often been performed in the absence of fluid flow, limiting the understanding of the effects of ubiquitous flow on TC transmigration and survival at the secondary site^4,19,20^. Studies that aim to recapitulate physiological fluid flows and shear stresses have so far been limited to the migration dynamics of TCs when embedded in 3D hydrogels^12,13,21^. For instance, Haessler *et al*. showed that interstitial fluid flow applied to MDA-MB-231 cells in matrigel enhanced their migration speeds, yet did not affect their directionality^12^. In another study, MDA-MB-231 cells embedded in a 3D collagen gel and subjected to physiological values of interstitial fluid flow^22,23^ were found to migrate in different directions depending on cell seeding density^13^. While these findings provide useful insight into the migration patterns of TCs generally in 3D matrices, they do not address the behavior of TCs in the vicinity of the vasculature and do not address the effects of local flow on TC extravasation and invasion.

Here, using *in vitro* microvascular networks (MVNs) that recapitulate human physiology^24^, a controlled hydrostatic pressure differential along and across the endothelium was applied to produce luminal and trans-endothelial flows of physiological magnitudes. By perfusing TCs in the vasculature, the individual and coupled effects of luminal and TE flows on both TCs and vasculature were quantified to understand their roles in extravasation and migration both intravascularly and in the surrounding matrix. While luminal flow was maintained in the microvasculature, TE flow was varied to recapitulate the range of physiological fluid flow profiles found in different vessels and tissues *in vivo*^7^. The results obtained help to elucidate the impact of these complex fluid flows on the metastatic potential of TCs.

## MATERIALS AND METHODS

### Cell culture and MVN formation

Pooled human umbilical vein endothelial cells (HUVECs, C2519A, Lonza) were transduced to stably express cytoplasmic fluorescence (LentiBriteTM GFP Control Lentiviral Biosensor 17-10387, Millipore Sigma) and cultured in Vasculife Endothelial Medium (LL-0003, Lifeline Cell Technology). Normal human lung fibroblast (NHLFs, CC-2512, Lonza) were cultured in Fibrolife Fibroblast Medium (LL-0011, Lifeline Cell Technology). Microfluidic devices (DAX-1, AIM Biotech) consisting of three three-channel chips on the same coverslip (central gel channel width of 1.3 mm and height of 250 μm) were used to form the MVNs. HUVECs were resuspended at 7 million cells/mL and NHLFs at 1 million cells/mL, both after seven passages, and seeded in fibrin hydrogel. A monolayer of HUVECs was added on the sides of the central gel channel on day 4 as previously described^9^, in this case to prevent direct migration of TCs into the matrix from the side channels. The MVNs were maintained in Vasculife Endothelial Medium, replenished every day for the duration of the culture. Devices were used between day 7 and day 12.

### TC perfusion and extravasation quantification

MDA-MB-231 immortalized breast cancer cells (HTB-26, ATCC) were cultured in Dulbecco’s Modified Eagle Medium (11995073, ThermoFisher) supplemented with 10% fetal bovine serum (16000044, ThermoFisher). TCs were resuspended in Vasculife Endothelial Medium at 1 million cells/mL and 10-20 μL of the suspension (100-200 cells) were perfused in the MVNs between days 7 and 10 under a transient flow *via* a hydrostatic pressure drop (by emptying the opposite media channel in each device). One microfluidic chip, containing three individual devices, was imaged at a time to quantify TC extravasation and migration. Devices were live-imaged on an Olympus FV1000 confocal microscope in a controlled environmental chamber (at 37 °C and 5% CO_2_)^20^ using a 10X objective and 5μm-thick z-stacks, choosing 2-5 regions of interest (ROIs) per device. TCs were perfused and maintained in the MVNs for 32 hours prior to the start of imaging, in order to ensure complete adhesion to the endothelium irrespective of flow condition imparted after that time. During the 32 hours, fewer than 5% of TCs had extravasated, significantly less than what was observed by our group with other batches of HUVECs, cultured in a separate channel from NHLFs^4,25^. In our current platform, HUVECs and NHLFs are cultured in the same gel channel which allows for both juxtacrine and paracrine signaling between cell types, along with a more physiological cellular organization. Co-culture with NHLFs is associated with improved vessel stability^26^ and barrier permeability (~ 20-fold reduction)^4,24,25^, a consequence of tighter endothelial junctions, and a possible reduction in the rate of TC extravasation compared to previous platforms where HUVECs and NHLFs were cultured in separate channels^4,25^. During imaging (16 hours), the effect of flow on TC extravasation and migration in the matrix was evaluated by quantifying extravasation efficiency of the TCs found to transmigrate during the imaging timeframe (removing from the analysis the TCs that extravasated in the first 32 hours), using the image analysis software Imaris (Bitplane) as previously described^4^. All TCs that were observed to partially or completely cross the endothelial barrier were classified as “extravasated”, and extravasation efficiency was computed as the ratio of extravasated TCs to total TCs (intravascular and extravasated) present in each fixed device. The rate of TC extravasation was measured using the live confocal images and defined as the fraction of extravasated TCs at any given time.

### Application and characterization of fluid flow

Luminal and TE flows were applied to the MVNs using two FlowEz pressure regulators (Fluigent), where tubing was connected to the devices using Luer connectors (LUC-1, AIM Biotech) as done previously^9^. In the fluidic system used (**Fig. 1A)**, two media channel outlets of the microfluidic chip on either side of the central gel channel were connected to media reservoirs pressurized by the FlowEz regulators, while the remaining two were closed. Luminal flow was produced across the MVNs through a pressure differential of 50 Pa between the side media channels, and it was coupled with TE flow by increasing the average media pressure to 500 or 1000 Pa compared to the open gel channel ports (atmospheric pressure). Luminal flow was characterized by measuring the velocity of fluorescently-labeled 2-μm diameter beads (F13083, Thermo-Fisher) in the MVNs as done previously^18^. The interstitial velocities resulting from TE flow in the matrix of the MVNs were quantified using a modified fluorescence recovery after photobleaching (FRAP) approach to track the movement of fluorescently-labeled 70kDa dextran within the interstitium, as done previously^9^. Four flow conditions were applied in this study: static control (no flow), 50 Pa luminal flow alone, 50 Pa luminal + 500 Pa TE (low-pressure trans-endothelial flow), and 50 Pa + 1000 Pa TE (high-pressure trans-endothelial flow) (**Fig. 1B**). For each chip containing three experimental sites, two were used as controls and one was subjected to a specified flow condition.

**Figure 1.**
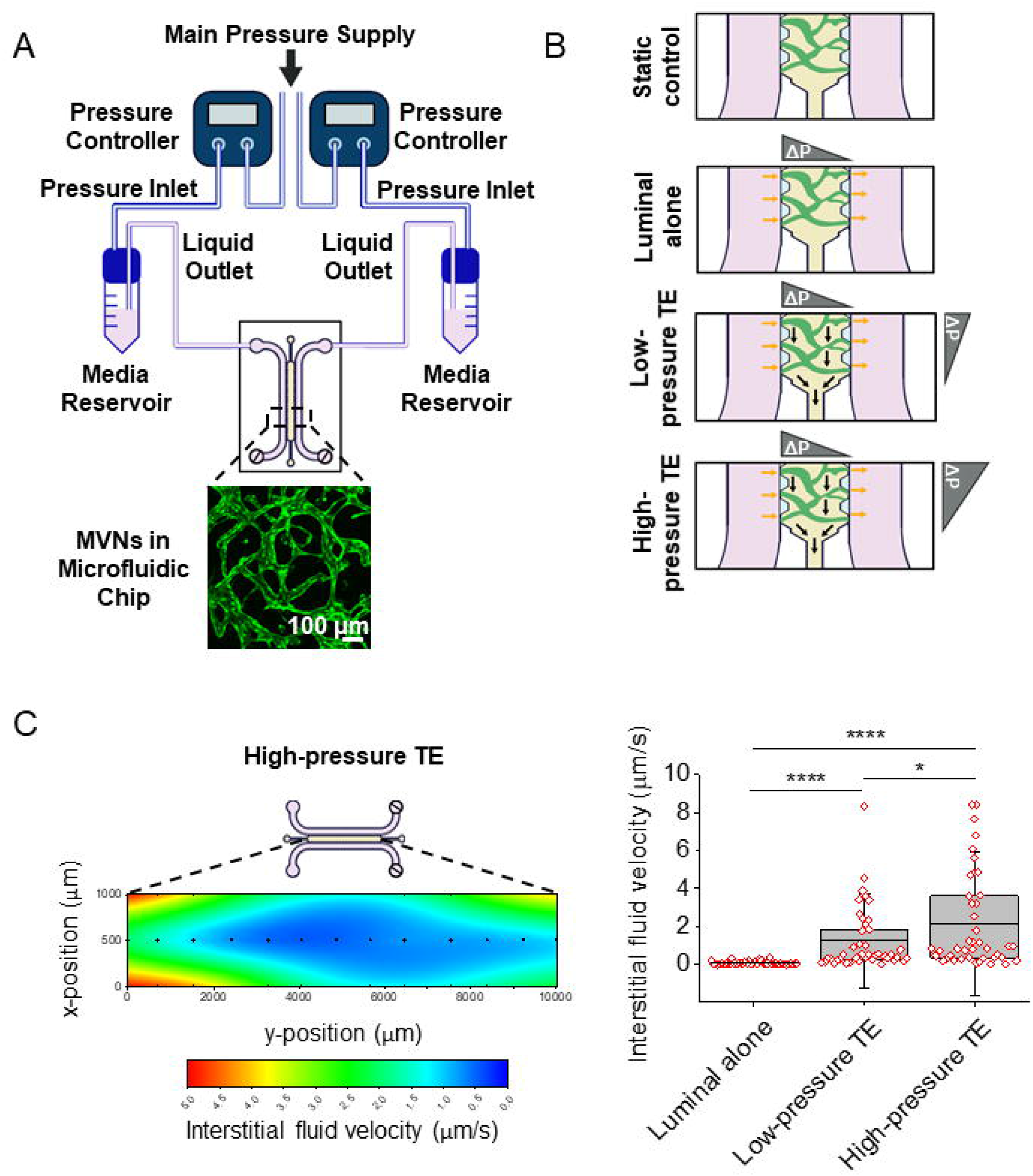
Flow characterization in human *in vitro* MVNs. **(A)** Diagram of the platform setup for the application of luminal and TE flows across perfusable MVNs (GFP HUVECs = green and non-fluorescent NHLFs) in a microfluidic chip connected to a pressure regulator. Both gel ports of the device are open, directing TE flow from the media channels through the MVN gel. **(B)** Schematic of the four different flow conditions applied to the MVNs. Control MVNs are not subjected to any flow, 50 Pa luminal are subjected to luminal flow across the gel of the microfluidic device, 50 Pa luminal + 500 Pa TE are subjected to both luminal and low-pressure TE flow from the media channels to the open gel ports, and 50 Pa luminal + 1000 Pa TE are subjected to both luminal and high-pressure TE flow. **(C)** Map of the interstitial fluid velocity distribution in the gel of the microfluidic device. Large flow profiles are observed near the liquid outlet ports of the media channels. **(D)** Box and whisker plot showing the distribution of interstitial fluid velocities in the gel under no flow (0 Pa), low-pressure TE flow (500 Pa) and high-pressure TE flow (1000 Pa). Mean and 25^th^ and 75^th^ percentiles are represented (n=42 ROIs per device for each flow condition).

### Cell migration analysis

In the 16-hour live-imaging timeframe, the trajectories and speed of migration of TCs were quantified via the Manual Tracking function of the TrackMate plugin in Fiji^27^. Briefly, the x-y coordinates of TCs were tracked at each time-point, and their speeds were obtained in μm/h from the tracking plugin. Although imaging was performed in 3D, the majority of the MVN vessel branches lie in the same z-coordinate plane owing to the small height of the microfluidic device (250 μm), and thus the speed and migration of TCs were computed in 2D. The x-y coordinates of TCs were adjusted for drift in 2D, which was quantified and corrected for in the analysis. TCs were classified as intravascular when travelling in the MVNs and extravascular after undergoing transendothelial migration and migrating in the matrix surrounding the MVNs. The intravascular or extravascular speed of TCs was measured as the average of all speeds for each pair of time-points prior to or following extravasation, respectively.

For TCs that eventually transmigrated, we analyzed their direction of motion pre- and post-extravasation. Pre-extravasation, cell migration direction was measured relative to the local luminal flow direction, assumed here to be aligned with the axis of the vessel branch in which they were located. The angles of extravasated TCs were measured relative to the y-axis corresponding to the predominant interstitial flow direction (parallel to gel channel). A linear regression was used to determine the axial orientation of each vessel branch using the coordinates of the endothelium, and consequently of luminal flow. The luminal flow direction obtained for each vessel branch was defined as the new x-axis, and TC coordinates were transformed to reflect this change. Using the adjusted x- and y-displacements of TCs, their angles θ with respect to the luminal flow direction were computed before extravasation, as shown in **SI Fig. 1**. All angles were grouped in 60-degree buckets between 0 and 360°, and graphed as angular displacement histograms using the PolarHistogram function of MATLAB (MathWorks) (**SI Appendix**). The migration distance of TCs with respect to the nearest endothelium during the 4 hours immediately following extravasation was also computed for all conditions.

### Finite-element computational model

To validate the fluid velocities in our microfluidic system and estimate the shear stress exerted on cells, a finite-element model was generated using COMSOL Multiphysics. A 3D geometry was used for the model with a single capillary, having the same diameter and endothelial hydraulic conductivity as previous studies from our group^24^, embedded in the central gel region, with a solid sphere attached to the luminal endothelial wall representing an adhered TC. Fluid transport was governed by Stokes and Brinkman-Darcy equations for fluid flow across the capillary and gel, respectively. Flow across the endothelium was governed by Starling’s Law and the hydraulic conductivity value corresponded to previous experimental measurements^9^. A steady flow was assumed, and the inlet/outlet boundaries were set to match the experimental conditions, a 50 Pa pressure difference for luminal flow and a 1000 Pa pressure difference for TE flow.

### Statistical analysis

All data are plotted as mean ± standard deviation, unless indicated otherwise. Statistical significance was assessed using student’s t-tests performed with the software OriginPro and represented as follows: n.s. stands for not significant, * denotes *p* < 0.05, ** denotes *p* < 0.01, *** denotes *p* < 0.001, and **** denotes *p* < 0.0001.

## RESULTS AND DISCUSSION

### Tunable fluid pressure differentials result in physiological values of luminal and trans-endothelial flows in the MVNs

To recapitulate physiological fluid flow in blood vessels and study its role in TC extravasation and invasion, we employed a previously developed microfluidic platform composed of perfusable MVNs in a fibrin gel^24,28^ (**Fig. 1A**). The application of a pressure differential between the two media channels generates luminal fluid flow that carries cell culture medium through the MVNs, as done previously^25^ (**Fig. 1B**). Simultaneously, TE flow can be generated by elevating luminal pressure relative to the surrounding matrix to drive fluid flow from the MVN lumens, between endothelial junctions^9^. The fluid leaving the MVNs enters the gel matrix, exiting via the open gel ports, to mimic interstitial flow and its drainage via the lymphatics^9^. The advantage of this approach lies in the fact that TE flow can be applied while maintaining the pressure differential necessary for luminal flow, thus recapitulating the presence of both flows in the microvasculature *in vivo* (**Fig. 1B**).

To characterize the different flow profiles within the system, MVNs were either subjected to luminal flow only, or luminal flow coupled with low- or high-pressure TE flow. Fluorescently-labeled beads, perfused in the MVNs under luminal flow resulting from a 50 Pa pressure difference between the two media channels, traveled with speeds up to ~ 500 μm/s, corresponding to a wall shear stress of up to ~ 0.15 Pa (**Movie 1, SI Fig. 2**). These flow speeds and corresponding shear stresses fall within the physiological ranges of 300 – 1500 μm/s and 0.05 – 0.25 Pa, respectively, observed in capillaries *in vivo*^15,29–32^, and are consistent with values previously used by our group in similar microfluidic platforms^18,25^. Interstitial fluid speeds in the matrix of the MVNs, resulting from the application of TE flow with larger pressure differentials (500 or 1000 Pa) across the endothelium, were quantified using fluorescence tracking. Increasing the pressure differential from 0 to 500 and 1000 Pa resulted in increased interstitial fluid velocities, from an average of 0.08 μm/s under 0 Pa, likely due to small and transient fluctuations in transendothelial pressure that caused minute interstitial flows, to values of 1.24 μm/s and 2.14 μm/s under 500 and 1000 Pa pressures, respectively (**Fig. 1C**). These interstitial fluid speeds fall within the range reported *in vivo* (0.1 – 4 μm/s)^22,23^. Both luminal and TE flows can be achieved simultaneously, thus enabling the study of TC extravasation and invasion in a highly physiologically relevant human-like microvasculature under appropriate flow conditions.

### Luminal and trans-endothelial flows differently affect tumor cell extravasation efficiency and rate

Using these physiologically relevant flows, we assessed their effect on TC extravasation. Breast MDA-MB-231 TCs were selected for this study given their well-characterized extravasation patterns in the MVNs and their studied migration patterns in 3D gels under interstitial flow^4,12,13,19^. Following perfusion of TCs under a transient flow and additional time to ensure TC adhesion, each microfluidic chip, either in static or flow conditions, was live-imaged under a confocal microscope for the entire duration of the experiment, *i.e*. 16 hours (**Fig. 2A, Movies 2-3, SI Movie 1**).

**Figure 2.**
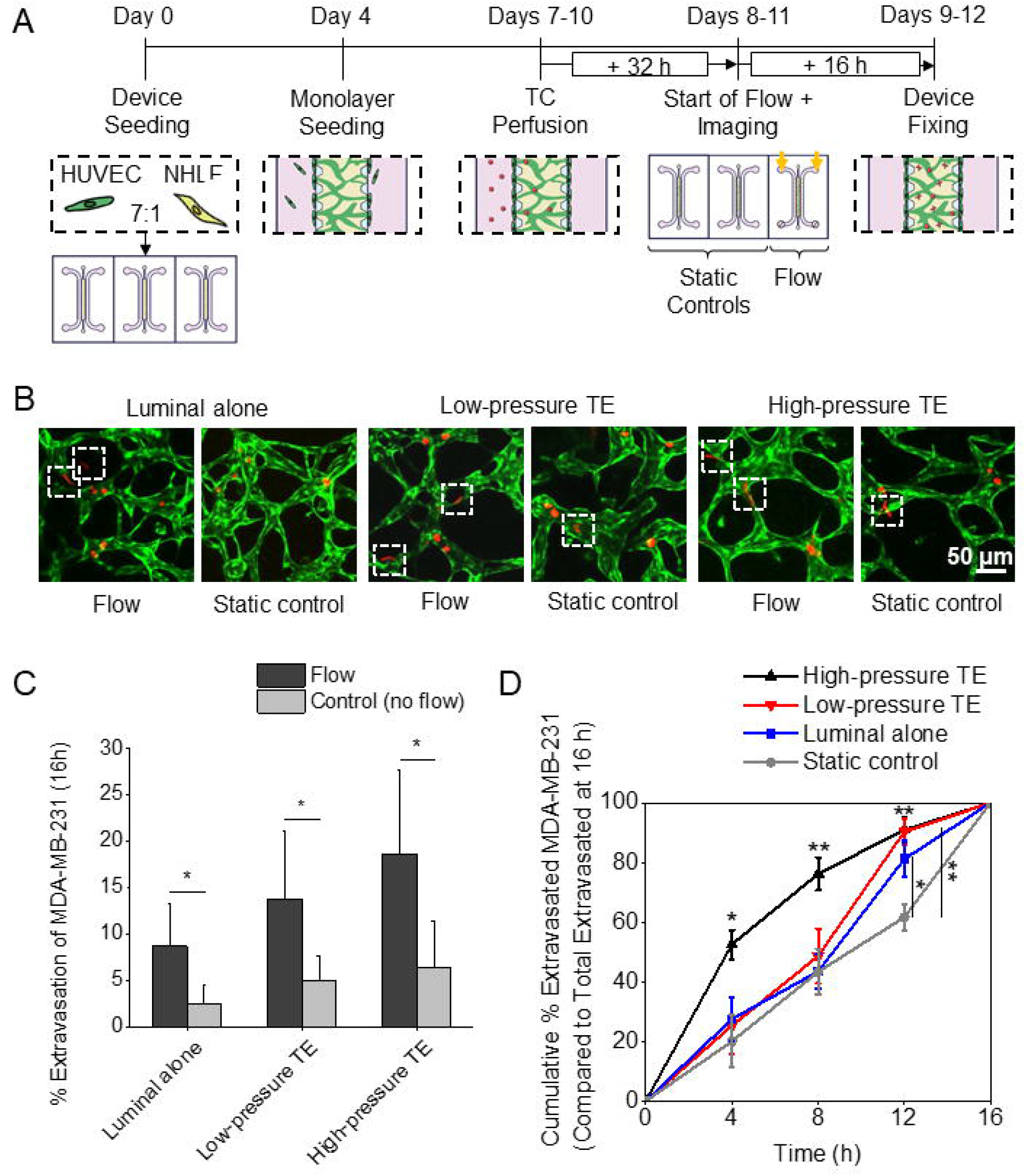
Effect of luminal and TE flows on the extravasation of TCs. **(A)** Schematic representing the sequential experimental protocol followed to obtain MVNs perfused with TCs and observe TC circulation, extravasation, and migration under flow. ~ 32 hours following TC perfusion, devices were subjected to flow and imaged under the confocal microscope for ~ 16 hours. **(B)** Representative collapsed z-stack confocal images of MVNs perfused with TCs under different flow conditions (and their respective controls without flow). Dashed square boxes indicate extravasated TCs (MDA-MB-231 cells = red, GFP HUVECs = green), recognizable by their elongated morphologies. All images were taken 6 hours following the start of flow. **(C)** Extravasation efficiency of MDA-MB-231 cells at the end of the confocal timelapses (t = 16 hour following start of flow) for all three flow conditions and their respective controls (n=6 devices per condition, 2-5 ROIs per device). **(D)** Extravasation rate of MDA-MB-231 cells corresponding to the cumulative percent of extravasated cells over the imaging timeframe (16 hours). Each flow condition is represented and all respective controls are pooled (n=6 devices per condition, 2-5 ROIs per device). Data is represented as mean ± standard error of the mean (S.E.M.), at the end of each four-hour imaging period. Significance is assessed with respect to control condition.

Extravasation of TCs from the MVNs was observed under all conditions investigated during the 16-hour imaging period (the small fraction (~5%) of TCs that extravasated in the first 32 hours was removed from the analysis to account only for the effect of flow on extravasation) (**Fig. 2B**). The timeline employed to assess extravasation in the MVNs (32-48 h) falls within the time needed for metastases to occur in animal models (1-3 days)^15,33^, thus supporting the relevance of our system in mimicking *in vivo* tumor progression. Using luminal flow alone or luminal flow coupled with TE flow resulted in significant increases in TC extravasation efficiencies, approximately 3-fold compared to static controls regardless of the type of flow applied (**Fig. 2C**). Indeed, when normalizing the values for each flow condition to their respective static controls, the presence of TE flow did not further alter TC extravasation efficiency (**SI Fig. 3**), suggesting that the larger number of TCs extravasating in all flow conditions was predominantly a result of luminal flow alone. TE flow did, however, affect the speed at which TCs extravasated from the MVNs. A significant delay in the extravasation of TCs can be observed in the case of low-pressure TE flow, luminal flow and static control, compared to high-pressure TE flow (**Fig. 2D**). Just four hours after the start of flow, more than half of all extravasated TCs had extravasated in the case of the high-pressure TE flow, compared to ~ 25% of all extravasated TCs for all other conditions, and this number rose to 75% halfway through the 16-hour imaging period, compared to ~ 45% for all other conditions (**Fig. 2D**).

This varied effect of luminal and transmural flow on TC extravasation is notable, as it suggests that TCs may possess a different extravasation potential depending on local flow conditions. It has previously been shown that endothelial cells are exquisitely sensitive to luminal shear stress and the resulting response may contribute to TC extravasation by changing endothelial morphology to facilitate the passage of TCs^15^. However, we did not observe any changes in endothelial cell morphology during TC extravasation under the different flow conditions tested, perhaps due to the relatively short period of flow. In addition, neither luminal nor transmural flows altered the micro-scale structure of the MVNs, as evidenced by a lack of variation in average vascular diameter over time between conditions (**SI Fig. 4**). These results, therefore, suggest that the changes in extravasation efficiency and rate stem primarily from alterations in the TCs under flow, as next investigated.

### Luminal flow, but not trans-endothelial flow, increases the intravascular migration speed of tumor cells

Following arrest and adhesion in the microcirculation, TCs show intravascular motility^5^, which was previously observed to also occur in the MVNs^4,18,25^. Using live confocal microscopy, the position of TCs in the microvessels was tracked to quantify migration speed (**Fig. 3A**). When comparing any type of flow with their respective controls, the average intravascular speed of TCs was found to be significantly higher with flow, increasing from ~ 9.4 μm/h to ~ 12.5 μm/h averaged across conditions (**Fig. 3B**). This was the case for luminal and low- and high-pressure TE flows and corresponded to an increase in overall distance travelled by the TCs in the MVNs; e.g.,~ 112 μm in a period of 7 hours prior to extravasation under high-pressure TE conditions compared to ~ 62 μm in the no-flow control (**Fig. 3A**).

**Figure 3.**
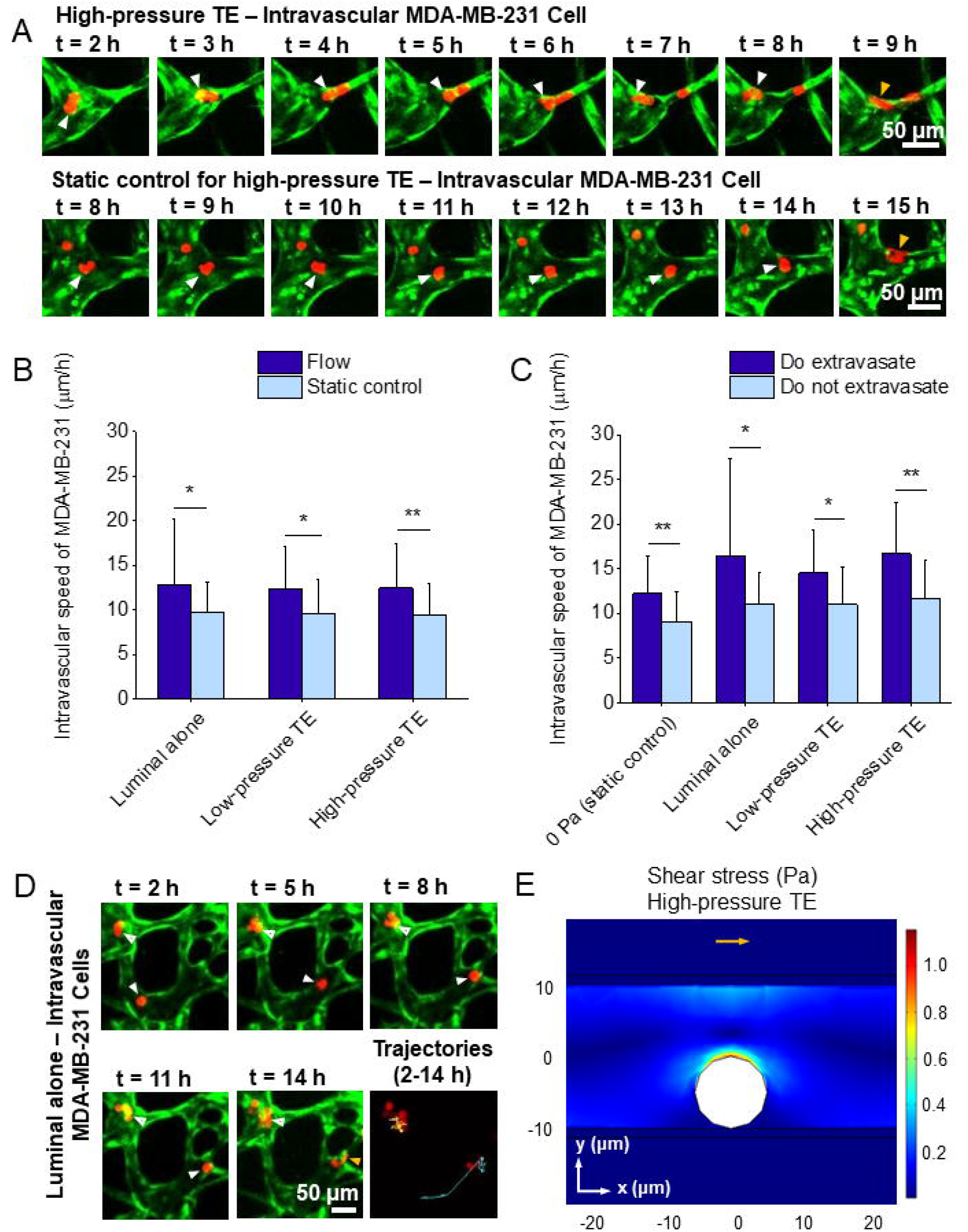
Effect of luminal and TE flows on the circulation of TCs in the MVNs. **(A)** Circulating MDA-MB-231 cells (red) in the MVNs (GFP HUVECs = green) before extravasation, under luminal and 1000 Pa TE flows (top) compared to its respective control MVNs without flow (bottom). The white arrows indicate the position of the TCs over time and the orange arrow in the last frame indicate the time of extravasation. The times correspond to the imaging intervals after the application of flow (total imaging time = 16 hours). **(B)** Intravascular speed of MDA-MB-231 cells before extravasation (in μm/h) for each flow condition and its respective control (n=6 devices per condition, 30-48 TCs considered per condition). **(C)** Intravascular speed of MDA-MB-231 cells that do and do not extravasate for each flow condition, including 0 Pa control devices without flow (n=6 devices per condition, 8-40 TCs considered per condition). **(D)** Circulating MDA-MB-231 cells (red) in the MVNs (GFP HUVECs = green) under 50 Pa luminal flow only. The open white arrows (top) indicate TCs that do not extravasate over the course of the imaging period and filled white arrows (bottom) indicate a TC that does extravasate at t = 14 hours (as shown by an orange arrow for extravasation). The intravascular trajectories of extravasated and non-extravasated TCs are shown (blue for extravasated cell at the bottom right and orange for non-extravasated cell at the top left). **(E)** Simulation of the shear stress profile on a TC adhered in a 3D microvessel under luminal and high-pressure TE flows, using a finite-element computational model. Luminal flow follows the direction indicated by the orange arrow and TE flow occurs in the perpendicular direction to the vessel.

The intravascular speeds of TCs were also compared for those that extravasate in the 16-hour imaging window as opposed to TCs that do not. This comparison was done to determine whether subpopulations of TCs might exhibit different migratory behaviors. Regardless of the type of flow applied, TCs that were observed to extravasate consistently exhibited increased intravascular speeds compared to TCs that did not (**Fig. 3C**). For instance, with luminal flow only, TCs that later extravasate were found to travel further intravascularly, compared to TCs in the same device that did not extravasate during imaging. As evidenced by their trajectories, TCs that extravasated 12 hours following the start of flow traveled an intravascular distance of ~ 200 μm prior to extravasation, while non-extravasating cells traveled a distance of ~ 50 μm intravascularly in the same time (**Fig. 3D**). Importantly, these results overall confirm the dominant role of luminal flow in determining the extravasation potential of TCs, irrespective of TE flow.

Shear stress from fluid flow has been found to increase TC migration and motility through mechanotransduction pathways^14,34,35^. Notably, focal adhesion protein re-organization has been observed in MDA-MB-231 cells and associated with their migration potential^14^. Yes-associated protein 1 (YAP1) was also found to be activated in prostate cancer cells subjected to fluid flow through activation of Rho kinase, which ultimately promoted their motility^34^. Here, the results suggest a potential mechanism through which luminal flow increases the migration speed of adhered TCs in the MVNs. In order to examine the role of luminal flow in producing shear stress on intravascular TCs, a computational model was developed to simulate luminal flow in conjunction with fluid loss through TE and interstitial flows (**Fig. 3E, SI Fig. 5**). The model was applied to a TC of diameter 10 μm adhered in a vessel of diameter 20 μm. Average luminal flow velocity near the TC was found to be ~ 4-times larger than far from the TC, resulting in a shear stress on the TC in the range of 0.3 – 1 Pa, sufficient to induce a mechanotransduction response^30,34,35^. This suggests that the increase in extravasation efficiency observed under flow in the MVNs could ensue from a shear stress-induced increase in cell migration due to activation of relevant transcription factors^34^. These shear stress levels could possibly result in an improved propensity to extravasate by facilitating the encounter of TCs with more permissive junctions between endothelial cells, as previously suggested^36^. Although shear stress has been found to decrease the permeability of endothelial barriers *in vitro*^19,37^, it has also been demonstrated that tight and adherens junction protein expressions are altered by the presence of physiologic levels of flow^38^. This reorganization of endothelial junctional proteins, along with flow-induced activation of transcription factors on TCs, could also influence extravasation efficiencies of TCs under flow.

### Trans-endothelial flow results in increased migratory speeds of extravasated TCs, which remain in close proximity with the endothelium

Flow-induced enhanced migration of TCs, as seen intravascularly under luminal flow, may also result from TE flow as this continues through the matrix as interstitial flow. Indeed, subjecting the system to rising interstitial fluid velocities (from 0.08 μm/s under luminal flow alone to a physiologically-relevant average of 2.14 μm/s under high-pressure TE flow) resulted in significant increases in the extravascular migration speed of TCs following extravasation (**Fig. 4A**). High-pressure TE flow resulted in a ~ 2.3-fold increase in the average extravascular speed of TCs in the matrix compared to static controls. Luminal flow alone, on the other hand, did not produce a similar increase. The effect of TE flow on TC extravascular migration speed also correlated with migration distances covered by the TCs, as shown in **Fig. 4B**, whereby TCs under high-pressure TE flow migrated an average distance of ~ 112 μm in the matrix during the 4 h after extravasation, while TCs under luminal flow alone traveled significantly less, only ~ 40 μm in the matrix during the same time period.

**Figure 4.**
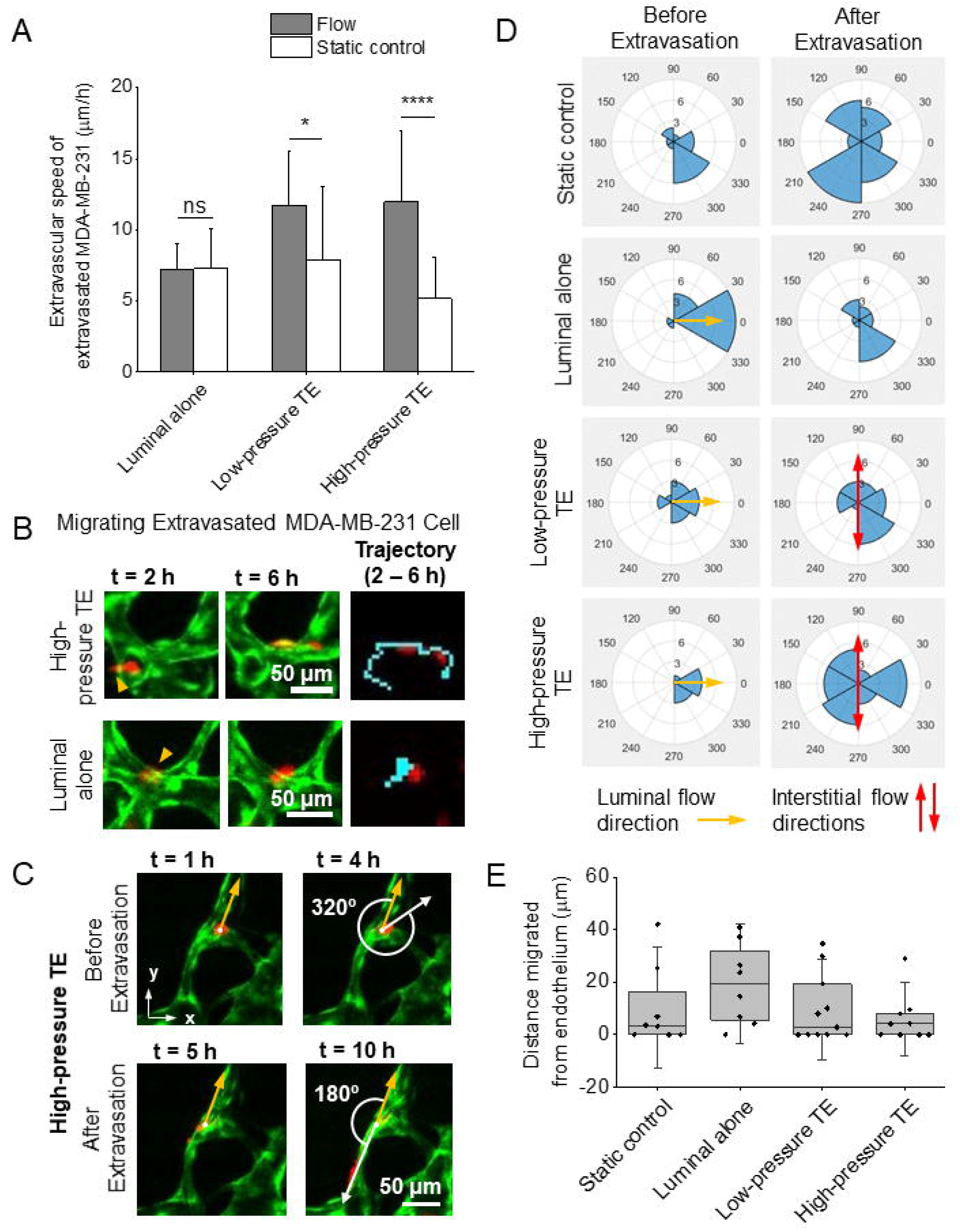
Effect of luminal and TE flows on the migration of TCs in the ECM. **(A)** Extravascular speed of MDA-MB-231 cells following extravasation and during migration in the ECM for each flow condition and its respective control (n=6 devices per condition, 5-28 TCs considered per condition). **(B)** Migrating MDA-MB-231 cells (red) following extravasation from the MVNs (GFP HUVECs = green) under luminal and 1000 Pa TE flows (top images) and luminal flow alone (bottom images). The orange arrow indicates the start of TC transmigration from the MVNs. Trajectories of migrating TCs in the extracellular space are shown in blue over the course of 4 hours following extravasation. **(C)** Representative images of the migration directionalities of an MDA-MB-231 cell (red) in MVNs (GFP HUVECs = green) under luminal and 1000 Pa TE flows. The orange arrow indicates the direction of luminal flow (along the vessel fragment considered). The TC is shown to circulate with an angle of 320° before extravasation (4 first hours) but adopts a migration angle of 180° in the matrix following extravasation (5 subsequent hours). **(D)** Polar histograms representing the distribution of TC migration angles in 60° buckets. Before extravasation, migration angles are quantified with respect to luminal flow in each vessel branch. After extravasation, migration angles of TCs are quantified with respect to the primary direction of interstitial flow (parallel to the gel channel). Quantifications are performed under different flow conditions (all controls are pooled here). The direction of luminal flow is represented by an orange arrow and corresponds to the direction of flow inside each vessel fragment considered for each TC. The direction of interstitial flow, following TE flow, is represented by the red arrows (parallel to the gel channel). **(E)** Distance migrated by TCs with respect to the nearest endothelium four hours following extravasation. All static controls were pooled from each flow condition. Mean and 25^th^ and 75^th^ percentiles are represented (n=6 devices per condition, 8-11 TCs considered per condition).

The role of flow on the directionality of motion of TCs, both intravascularly and in the ECM following extravasation, was assessed next. The directional bias of TCs before extravasation was quantified by measuring the angle adopted by TCs with respect to the direction of luminal flow in their individual vessel fragments and classifying it into polar histograms with 60° buckets (**Fig. 4C, SI Fig. 1**). The directional bias of extravasated TCs was also quantified with respect to the primary direction of interstitial flow (parallel to the gel channel). For any type of flow applied in the MVNs, adherent TCs were observed to travel predominantly in the direction of luminal flow before extravasation (**Fig. 4D**). For example, nearly 70% of intravascular TCs under both luminal and high-pressure TE flow travelled in the same direction as that of luminal flow before extravasating (330-30° angle bucket). As expected, in the static controls there was no directional preference. This is consistent either with a mechanotransducive role of luminal flow on TCs, or simply a physical one, in which the shear stress drags the cell in the flow direction.

After extravasation, TCs adopted a very different pattern of directional migration (**Fig. 4D**). Strikingly, the addition of TE flow did not result in any specific migration directionality for extravasated TCs, when compared to the static and luminal flow conditions. This could be due to the presence of large pressure differences in the TE flow conditions which generate more complex interstitial flow profiles, resulting in extravasated TCs that span all angular directions. Overall, these findings show that TE flow results in increased interstitial fluid velocities promoting the migration of extravasated TCs, which has been linked to improved survival and successful colonization of the metastatic niche^39,40^. Moreover, TCs under TE flow-derived interstitial flow migrated in close proximity to the vasculature following extravasation in the ECM. This was not the case for TCs under luminal flow alone, which migrated further away from the endothelium on average, possibly because of the uni-directional interstitial flow (**Fig. 4E**). Close association with the vasculature following transmigration, also referred to as co-option^41^, has been linked with optimal oxygen supply and access to nutrients for TCs, ultimately leading to TC survival and proliferation^42^. Indeed, *in vivo* observations of metastatic colonization suggest that vessel co-option, which has been reported in lung metastases from patients with primary breast cancer^41^, is a key determinant of the overall survival of TCs in the matrix following extravasation, and governs the formation of micrometastasis^33^. In some experiments, extravasated TCs under high-pressure TE flow were observed to undergo cell division when migrating in the ECM close to the endothelium (**Movie 4, SI Movie 2**). These observations suggest a possible role for TE flow in promoting TC proliferation following extravasation, another key feature of the formation of successful metastatic foci^33^. Finally, the enhanced TC extravasation rates under high-pressure TE flow (**Fig. 2D**) further suggest that TE flow may even directly affect TCs during trans-endothelial migration by increasing their migration speed across endothelial junctions, overall enhancing the invasiveness of TCs from the moment they begin leaving the microvasculature.

## CONCLUSIONS

These studies identify an important role for luminal and trans-endothelial flows in promoting tumor cell extravasation and migration in the surrounding matrix, *via* the action of physiological fluid flow on both TCs and MVNs. In particular, our results show that intravascular flow can promote tumor cell extravasation, despite previously observed reductions in barrier permeability under flow^19,25^. Flow-induced endothelial junctional reorganization and elicited mechanotransduction responses in TCs under increased shear stresses can therefore play a major role in promoting TC extravasation. This result attests to the importance of applying physiological luminal flow profiles when studying extravasation *in vitro*, in addition to the use of models that can capture human pathophysiological phenomena with high spatio-temporal resolution^43^. TE flow was found to increase both the speed at which TCs transmigrate as well as their migratory speeds once in the surrounding matrix. These mechanisms are consistent with previous observations in an *in vitro* system of the lymphatic vasculature, in which TE flow increased transmigration rates of TCs through an upregulation of key ligands and receptors involved in TC transmigration^44^. Such upregulation may provide a lead for future work to further understand the effect of TE flow on TCs. Here, extravasated TCs under TE flow were also found to remain close to the vasculature, further emphasizing the role of TE flow in promoting the successful formation of metastatic foci. Overall, the studies conducted offer valuable insights into the effects of flow on extravasation and migration, suggesting a role for local flow rates in determining the metastatic potential of TCs. Future work should more broadly map these effects on different TCs exhibiting divergent migration patterns^45^. Ultimately, these studies may lead to new therapeutic strategies targeted to specific cancer cells/sites in the context of fluid flows present at these locations.

## AUTHOR CONTRIBUTIONS

C.H., L.I., J.C.S., G.S.O., and R.D.K. designed the research; C.H., L.I., J.C.S., and G.S.O. performed the experiments; C.H. and L.I. analyzed the data pertaining to tumor cells; G.S.O. analyzed the data pertaining to flow characterization; J.C.S developed the finite-element computational model; C.H. wrote the first draft of the manuscript, all authors contributed to its final form.

## ACKNOWLEDGEMENTS

C.H. is supported by the Ludwig Center for Molecular Oncology Graduate Fellowship and by the National Cancer Institute (U01 CA202177). L.H. is supported by the National Cancer Institute (U01 CA202177). J.C.S. is supported by an NSF Graduate Research Fellowship. G.S.O. is supported by an American-Italian Cancer Foundation Post-Doctoral Research Fellowship. This work was supported by a National Cancer Institute Physical Sciences-Oncology Network supplement to award U01 CA202177. The authors thank Dr. Mark Gillrie for his help transfecting HUVECs with the GFP construct.

---

**Movie 1.** Movie showing beads flowing in the MVNs under luminal flow (beads = pink, GFP HUVECs = green). Images taken every 0.09 seconds.

**Movie 2.** MDA-MB-231 cells (red) circulating, extravasating, and migrating in the MVNs (GFP HUVECs = green) under luminal and 1000 Pa TE flows, over the course of 16 hours. Images taken every hour.

**Movie 3.** MDA-MB-231 cells (red) circulating, extravasating, and migrating in the MVNs (GFP HUVECs = green) without any flow (control), over the course of 16 hours. This movie is the respective control of Movie 2 under luminal and 1000 Pa TE flows. Images taken every hour.

**Movie 4.** MDA-MB-231 cell (red) extravasating, migrating, and subsequently undergoing mitosis in the MVNs (GFP HUVECs = green) under luminal and 1000 Pa TE flows. Images taken every hour.

## DATA AVAILABILITY

The research data required to reproduce these findings is available upon request from the corresponding author.

