## Supplementary figures and images for "The effects of luminal and trans-endothelial fluid flows on the extravasation and tissue invasion of tumor cells in a 3D *in vitro* microvascular platform"

### Supplemental Figure 1

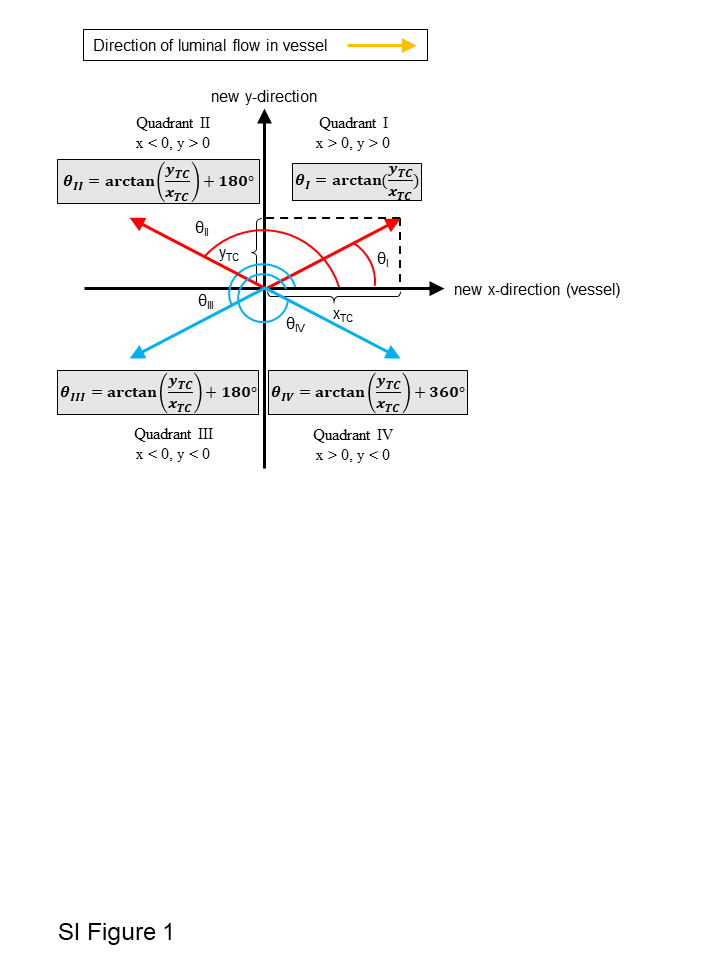

### Supplemental Figure 2

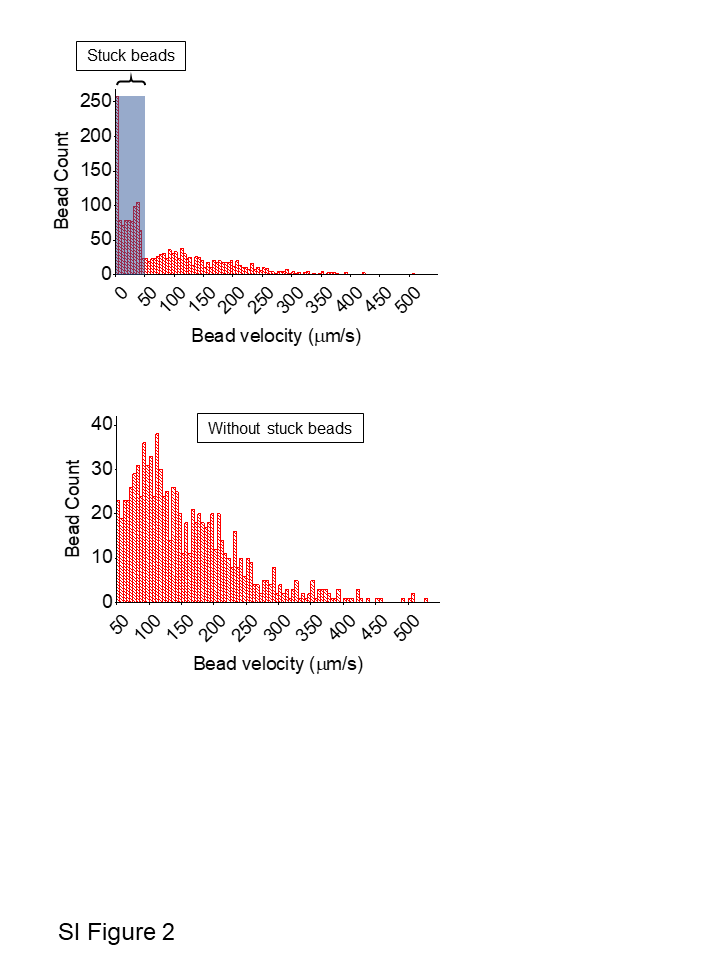

### Supplemental Figure 3

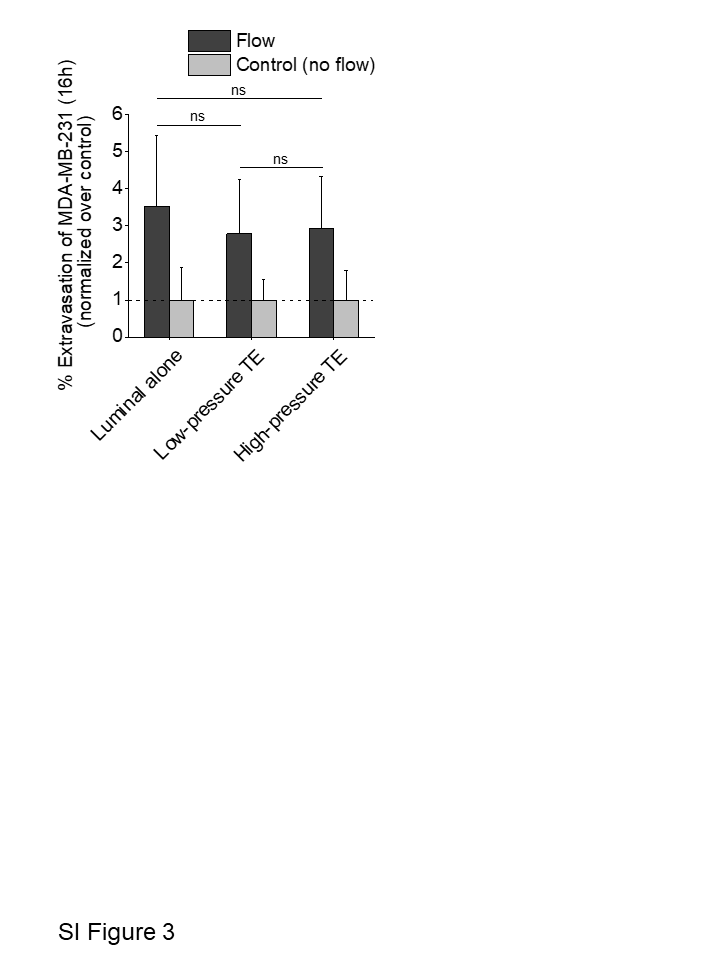

### Supplemental Figure 4

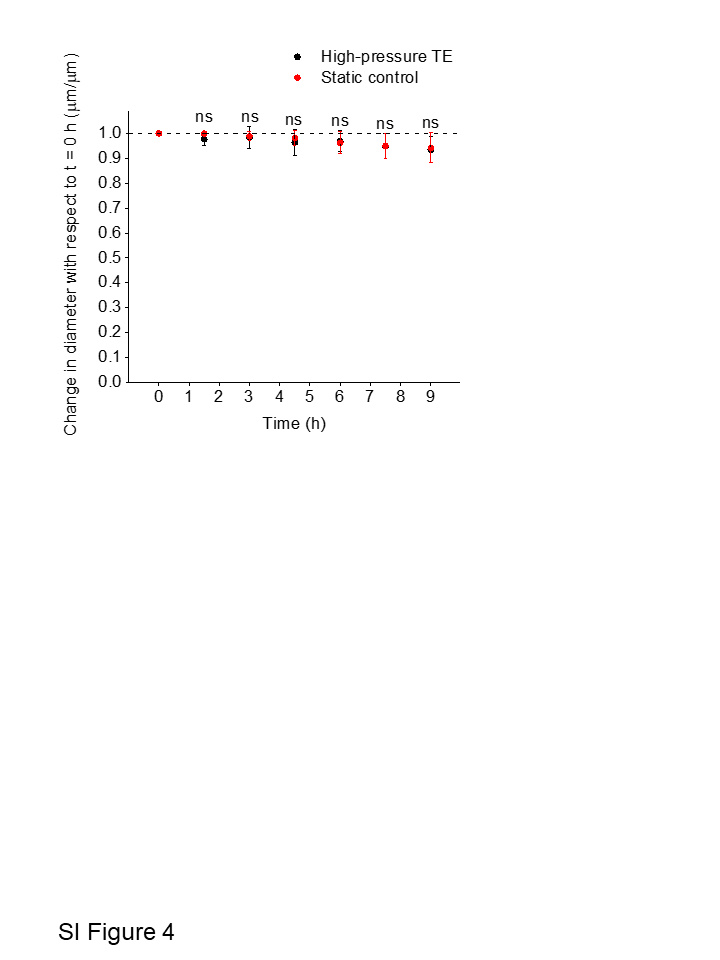

### Supplemental Figure 5

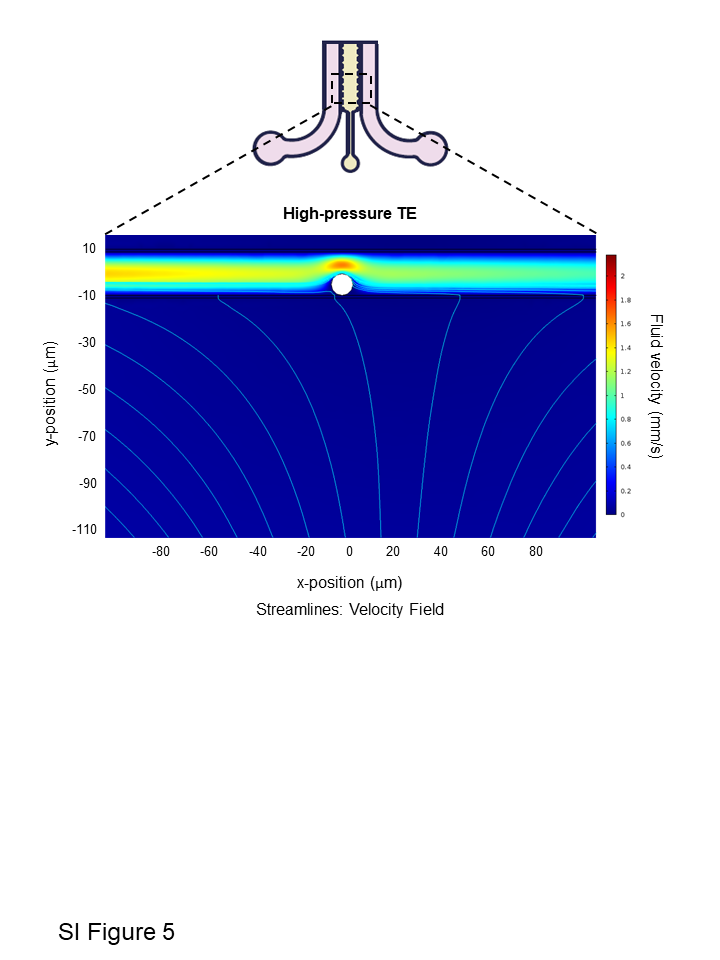
