## Supplemental Information for "The effects of luminal and trans-endothelial fluid flows on the extravasation and tissue invasion of tumor cells in a 3D *in vitro* microvascular platform"

**SI FIGURE/MOVIE LEGENDS**

**SI Figure 1. Schematic for the measurements of angular displacements of TCs (with respect to the luminal flow direction in each vessel considered).** TC coordinates are adjusted to ensure that its x-motion is in the same direction as that of the vessel and consequently luminal flow. The angle θ can be computed using an arctangent formula depending on the quadrant the TC is located in. The direction of luminal flow is indicated by an orange arrow.

**SI Figure 2. Graph of bead count per luminal bead velocity. *Top.*** Graph of the distribution of beads with respect to luminal velocity showing that a large fraction of beads remains stuck in the MVNs with speeds approaching zero (n=1,858 beads considered). ***Bottom.*** Adjusted graph of the distribution of “moving” beads with respect to luminal velocity (n=737 beads considered).

**SI Figure 3. Extravasation efficiency of MDA-MB-231 cells at the end of the confocal timelapses (16 h) normalized with respect to respective controls for each type of flow.** Extravasation efficiencies of each flow condition are normalized to control extravasation efficiencies. TE flow does not improve extravasation efficiencies compared to luminal flow (n=6 devices per condition, 2-5 ROIs per device).

**SI Figure 4. MVN morphology comparison under luminal and TE flow compared to control (no flow).** Ratios of the average MVN diameter at each timepoint compared to the initial average diameter, for conditions of luminal and 1000 Pa TE flow compared to control (no flow). Average diameters were computed over the course of 9 hours under flow or no flow at 1.5 hour intervals. Changes in diameters are not significantly different from one another under flow or no flow (n=4 devices per condition).

**SI Figure 5. Simulation of the fluid velocity profile on a TC adhered in a 3D microvessel under luminal and 1000 Pa TE flows.** Fluid velocity profile was mapped in an ROI near the open gel port of the central region of the microfluidic device (velocity streamlines are indicated in the vessel and matrix). Increasing fluid velocities are observed near the TC in the vasculature.

**SI Movie 1.** MDA-MB-231 cells (red) circulating, extravasating, and migrating in the MVNs (GFP HUVECs = green) under luminal flow alone, over the course of 16 hours. Images taken every ~ 30 min.

**SI Movie 2.** MDA-MB-231 cell (red) extravasating, migrating, and subsequently undergoing mitosis in the MVNs (GFP HUVECs = green) under luminal and 1000 Pa TE flows. Images taken every ~ 30 min.

**SI APPENDIX**

clear

close all

fprintf('Loading Data from Excel\n');

angles1000_cutoff = xlsread('MigrationDataAnglesMatlabBeforeExtrav.xlsx','AK3:AK47');

angles500 = xlsread('MigrationDataAnglesMatlabBeforeExtrav.xlsx','AN3:AN37');

anglesluminal = xlsread('MigrationDataAnglesMatlabBeforeExtrav.xlsx','AQ3:AQ44');

anglescontrol_cutoff = xlsread('MigrationDataAnglesMatlabBeforeExtrav.xlsx','AT3:AT87');

fprintf('Convert to Radians\n');

angles1000_cutoff = angles1000_cutoff*pi/180;

anglescontrol_cutoff = anglescontrol_cutoff*pi/180;

angles500 = angles500*pi/180;

anglesluminal = anglesluminal*pi/180;

bin_number = 6;

bins = linspace((pi/6),((13*pi)/6),bin_number+1);

subplot(1,4,1)

polarhistogram(angles1000_cutoff,'BinEdges',bins)

title('1000 Pa');

rlim([0 9])

rticks([3 6])

subplot(1,4,2)

polarhistogram(angles500,'BinEdges',bins)

title('500 Pa');

rlim([0 9])

rticks([3 6])

subplot(1,4,3)

polarhistogram(anglesluminal,'BinEdges',bins)

title('Luminal');

rlim([0 9])

rticks([3 6])

subplot(1,4,4)

polarhistogram(anglescontrol_cutoff,'BinEdges',bins)

title('Control');

rlim([0 9])

rticks([3 6])
